# The 2.3 Å Structure of A21, a Protein Component of the Conserved Poxvirus Entry-Fusion Complex

**DOI:** 10.1101/2025.01.08.631918

**Authors:** Ulrike S. Diesterbeck, Liya A. Muslinkina, Apostolos G. Gittis, Kavita Singh, Bernard Moss, David N. Garboczi

## Abstract

Poxviruses are exceptional in having an entry-fusion complex (EFC) consisting of eleven conserved proteins embedded in the membrane of mature virions. With the goal of understanding the function of the EFC, extensive efforts have been made to determine the structures and roles of its components, and to date, structures have been determined for nine of the eleven proteins. Here, we report the crystal structure of A21, the 10^th^ EFC protein, comprising two α-helices clasping a twisted antiparallel β-sheet stabilized by two conserved disulfide bonds. The stability of each of the three A21 loops is provided by hydrogen bonds between main-chain atoms and several highly conserved residues, making the overall fold of A21 and its orthologs resilient to evolutionary change. Based on AlphaFold modeling and phylogenetic analysis of A21, we suggest that its highly conserved N-terminal transmembrane domain and C-terminal α-helix enable A21 integration into EFC, where it primarily interacts with the G3/L5 subcomplex and the smallest of EFC components, the O3 protein.

**Highlights:**

- With the structure of A21 determined by us, the structures of 10 out of the 11 proteins forming the entry-fusion complex (EFC) of the poxvirus are now known.
- A21 features a highly conserved transmembrane (TM) domain, which, along with the C-α-helix, facilitates the integration of A21 and its orthologs into the EFC.
- Within the EFC, A21 primarily interacts with the G3/L5 subcomplex and the O3 peptide.

## Introduction

The poxviruses comprise a large family of enveloped double-stranded DNA viruses that infect many vertebrates and invertebrates and are responsible for human diseases including smallpox and molluscum contagiosum as well as mpox and other zoonoses [1]. In particular, the increasing incidence of mpox in Africa and the global spread due to human-to-human transmission have motivated efforts to better understand the biology of poxviruses and develop more effective vaccines and therapeutics [2]. Unlike other DNA viruses, poxviruses replicate entirely in the cytoplasm and encode their own unique transcription and replication systems [3]. Another key difference from other viruses is the poxvirus mode of cell entry, which can occur by fusion with the plasma or endosomal membrane. Whereas most enveloped viruses have one or two proteins devoted to membrane fusion, poxviruses have 11 small proteins that form the entry-fusion complex (EFC) and are each required for infection [4]. Orthologs of each EFC protein are highly conserved and present in nearly all poxviruses including those that infect vertebrates and invertebrates. Moreover, some EFC homologs are also present in African Swine Fever Virus, a member of the Asfarviridae family [5] and giant viruses belonging to the Nucleocytoviricota [6]. However, only the entry-fusion proteins expressed by vaccinia virus (VACV), the prototype poxvirus, have been characterized experimentally and demonstrated to exist in a complex. VACV EFC proteins A16, G9 and J5 have C-terminal transmembrane (TM) domains, conserved intramolecular disulfide bonds and partial sequence identity indicating an origin from a common ancestral gene [7–9]. L1 and F9 have C-terminal TM domains, partial sequence identity and likely also arose by gene duplication [10, 11]. The other EFC proteins are unrelated: L5 has a C-terminal TM domain [12], whereas A21, A28, G3, H2, and O3 have N-terminal TM domains [13–15]. None of the EFC proteins have cleavable signal peptides or are glycosylated as they apparently bypass the endoplasmic reticulum to insert directly in the cytoplasmic viral membrane [16]. Another unique feature of the EFC proteins is the presence of intramolecular disulfide bonds that are formed by a highly conserved poxvirus cytoplasmic redox system [17]. Entry entails a hemifusion step, in which lipid mixing of viral and cellular membranes occurs, followed by pore formation and penetration of the core into the cytoplasm [18]. However, neither a putative cell activation receptor nor the mechanism of fusion has been determined to date.

Elucidation of the structure of the EFC may help to understand the fusion process and point to improved therapeutics and vaccines. Although the overall structure of the EFC has not been determined, several protein-protein interactions have been found including A28:H2 [19], A16:G9 [20], and G3:L5 [21]. At present, structures have been reported for the ectodomains of nine out of the eleven proteins forming the EFC: L1 [22], F9 [23], the heterodimers of G3:L5 [24] and A16:G9 [25], A28 [26], H2 [27], and J5 [28].

The present study examines the structure of VACV A21 (OPG147), one of the two remaining uncharacterized EFC proteins. A21 synthesis occurs at a late stage of virus infection and the protein localizes on the lipoprotein membrane of intracellular mature virions (MVs) [29]. Studies with a conditional lethal mutant in which the A21 gene was down regulated by the lac repressor demonstrated that virions lacking A21 were able to bind to cells, but were unable to mediate cell fusion [29]. The association of A21 with the EFC was demonstrated by affinity chromatography and mass spectrometry [9]. Here we report on the experimentally determined structure of the ectodomain of A21, its conservation in other poxviruses, and predicted interactions with the other proteins of the poxvirus EFC.

## Results and Discussion

### The structure and topology of the poxvirus A21 protein

We expressed the ectodomain of the VACV strain Western Reserve (WR) A21, consisting of amino acid residues 24–117 with a six-His tag and protease cleavage site at the N-terminus (Fig. 1A). The protein was refolded from inclusion bodies (IBs) produced in bacteria and purified to near homogeneity by immobilized metal affinity chromatography and size exclusion chromatography (SEC). The integrity of recombinant A21 was verified by dynamic light scattering (DLS), which showed a low polydispersity consistent with a uniformly monomeric protein sample. Rod-shaped crystals with a hexagonal cross-section were obtained by hanging drop vapor diffusion.

**Figure 1.**
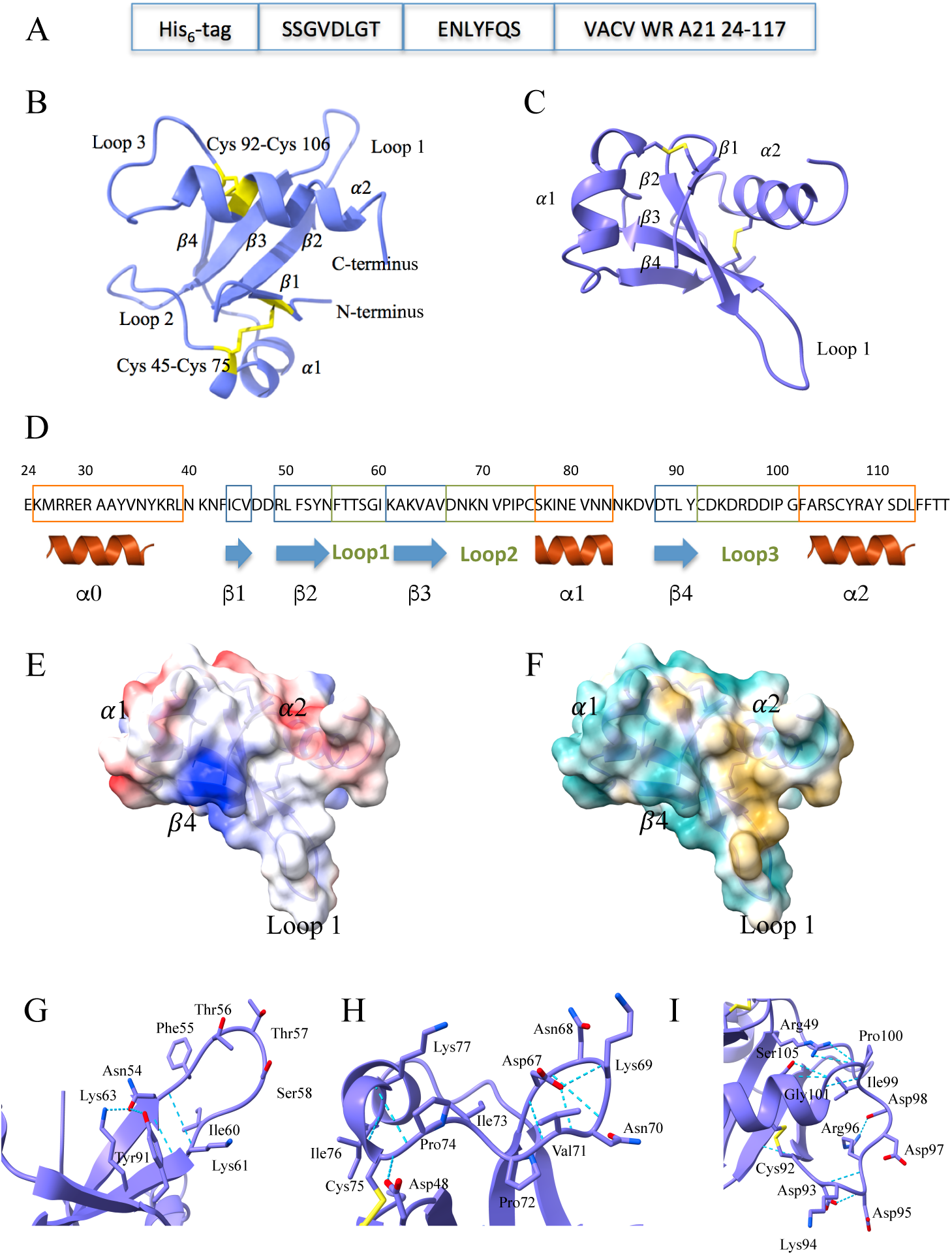
A21 structure and analysis. **A**, Expression construct of the A21 ectodomain. Shown are the His_6_-tag for affinity purification, the SSGVDLGT linker, the tobacco etch virus (TEV) protease recognition sequence ENLYFQS, and residues 24-117 of A21 from vaccinia virus Western Reserve strain (VACV WR). **B**, X-ray crystal structure of A21. Ribbon trace (purple) and disulfide bonds (yellow) are shown together with the two α-helices (α1-α2), four β-strands (β1: Ile44-Val46, β2: Arg49-Asn54, β3: Lys61-Val66, and β4: Asp88-Tyr91), and three loops (Loop 1-Loop 3). The two disulfide bonds are labeled. **C**, X-ray crystal structure of A21 rotated and translated from panel B. **D**, Electrostatic potential surface of A21 shown in the same orientation as in panel C and based on the X-ray structure (residues 41-117). The color varies from more negative surface potential (red) to zero potential (white) to more positive potential (blue). **E**, Hydrophobicity surface of A21 in the same orientation as in panel C. The color varies from a more hydrophobic surface (yellow) to a more hydrophilic surface (cyan). **F,** Close-up of Loop 1 (Asn54-Ile60) and the hydrogen bonds (blue dashed lines) that stabilize it. **G**, Close-up of Loop 2 (Asp67-Cys75) and its hydrogen bonds. **H**, Close-up of Loop 3 (Cys92-Gly101) and its hydrogen bonds. **I**, Secondary structure diagram of the A21 ectodomain. The α0-helix is shown as predicted from the residues 24-39, which though expressed, were not seen in the electron density.

We determined the structure of A21 to 2.3 Å resolution by X-ray crystallography using phase estimates calculated from data collected using 1.54 Å wavelength X-rays from A21 crystals soaked in sodium iodide (Supplementary Table 1). The asymmetric unit contains six A21 molecules (A to F) stabilized by interactions along two kinds of interfaces (Fig. S1). The first kind are the identical A/B, C/F, and D/E interfaces, which consist of antiparallel β-sheets formed by the β4-strands from each subunit. The A/B interface has an area of about 650 Å^2^ and is stabilized by four hydrogen bonds. The second type of interface in the asymmetric unit is between A/D and B/C with an area of about 690 Å^2^. The interface is stabilized by two hydrogen bonds and by hydrophobic interactions between loop 1 and the α2-helix of each molecule (Fig. S1).

The structure of the A21 ectodomain comprises two α-helices clasping a bundle of four β-strands that form a twisted antiparallel β-sheet (Fig. 1B-C). The disulfide bond Cys45-Cys75 stabilizes the spatial arrangement of β1-strand and α1-helix; the disulfide bond Cys92-Cys106 stabilizes the positioning of α2-helix, β4-strand, and the base of loop 3. The A21 ectodomain has two negatively charged clusters at loop 3 (Asp93, Asp95, Asp97, Asp98) and located at the residues connecting the α1-helix and β4 strand (Glu80, Asp86, Asp88) (Fig. 1E). Lys61, Lys63, and Lys85 form a large positively charged patch (Fig. 1E). A21 has an overall hydrophilic surface (cyan, Fig. 1F) and a hydrophobic area (yellow) formed by the residues of the α2-helix, β3-strand, and loop 1: Ile44, Val46, Phe51, Tyr53, Phe55, Ile60, Ala109, Tyr110, and Leu113. Smaller hydrophobic patches are comprised of Ile72, Ile78, Val81, Leu90, Tyr91, Phe114 and Phe115 (Fig. 1E). The charged residues and hydrophobic patches are possible sites for A21 interactions with the other EFC proteins.

Additional structural elements with a potential for binding to the components of the EFC are loops 1-3 (Fig. 1 F-I). Loop 1 is stabilized at its base by a pair of hydrogen bonds between Asn54 and Lys61 (Fig. 1F). It also participates in the formation of the A/D interface, where loops 1 of the molecules A and D are arranged in an antiparallel fashion (Fig. S1A, purple and beige). The shape of loop 2 is stabilized by hydrogen bonds (Fig. 1G). The side chain of Asp67 extends inside loop 2 and forms hydrogen bonds with the main chain nitrogen atoms of Lys69 and Val71. Additionally, the main chain oxygen and nitrogen of Asp67 participate in the hydrogen bonding with the main chain Lys69(N) and Val71(O). The side chains of the loop residues Asn68, Lys69, and Asn70 point outwards and do not form hydrogen bonds. Two other main chain hydrogen bonds with residues outside the loop, between Ala65 and Ile73 and between Pro74 and Lys77, further stabilize the position of loop 2 relative to the β3-strand and α1-helix (Fig. 1G).

Loop 3 bears numerous amino acid residues with negatively charged side chains (Fig. 1H). The shape of loop 3 is stabilized by hydrogen bonds between the main chain atoms of Asp93 and Arg96, Ile99 and Phe102, and Gly101 and Ser105. The hydrogen bonding between the side chains of Asp93 and Asp95 and between Arg96 and Asp98 further restricts the mobility of loop 3 and defines its shape. Additional hydrogen bonds between the residues of loop 3, β2- and β3-strands, and α2-helix residues stabilize its position relative to the rest of A21: Ala62(O) and Cys92(N), Arg49(NH1) and Pro100(O), Arg49(NH2) and Pro100(O), Ser105(OG) and Gly101(O). Thus, loops 2 and 3 with charged amino acid residues facing outwards present potential sites for association with the other proteins of the EFC. Extensive hydrogen bonds between main chain atoms in loops 2 and 3 may provide evolutionary stability of their shapes, as they are independent of the side chains of the respective residues.

The A21 ectodomain structure predicted with AlphaFold 2.0 using the sequence that was expressed (24-117) agreed with our X-ray crystal structure, and also suggested that the first 18 amino acid residues (residues 24-41) form a large α0−helix (cyan helix, Fig. 2A). We do not see the α0−helix in our crystal structure presumably due to its flexible connection to the rest of the molecule in the absence of the N-terminal TM segment. Modeling with the full-length sequence (1-117) revealed that the α0− and transmembrane helices are predicted to be almost continuous (Fig. 2B and Fig. S2). The α0−helix changes the electrostatic and hydrophobic surfaces of the molecule, by presenting a new positively-charged surface and covering the hydrophobic area at the base of loop 1 (Fig. 2C-F).

**Figure 2.**
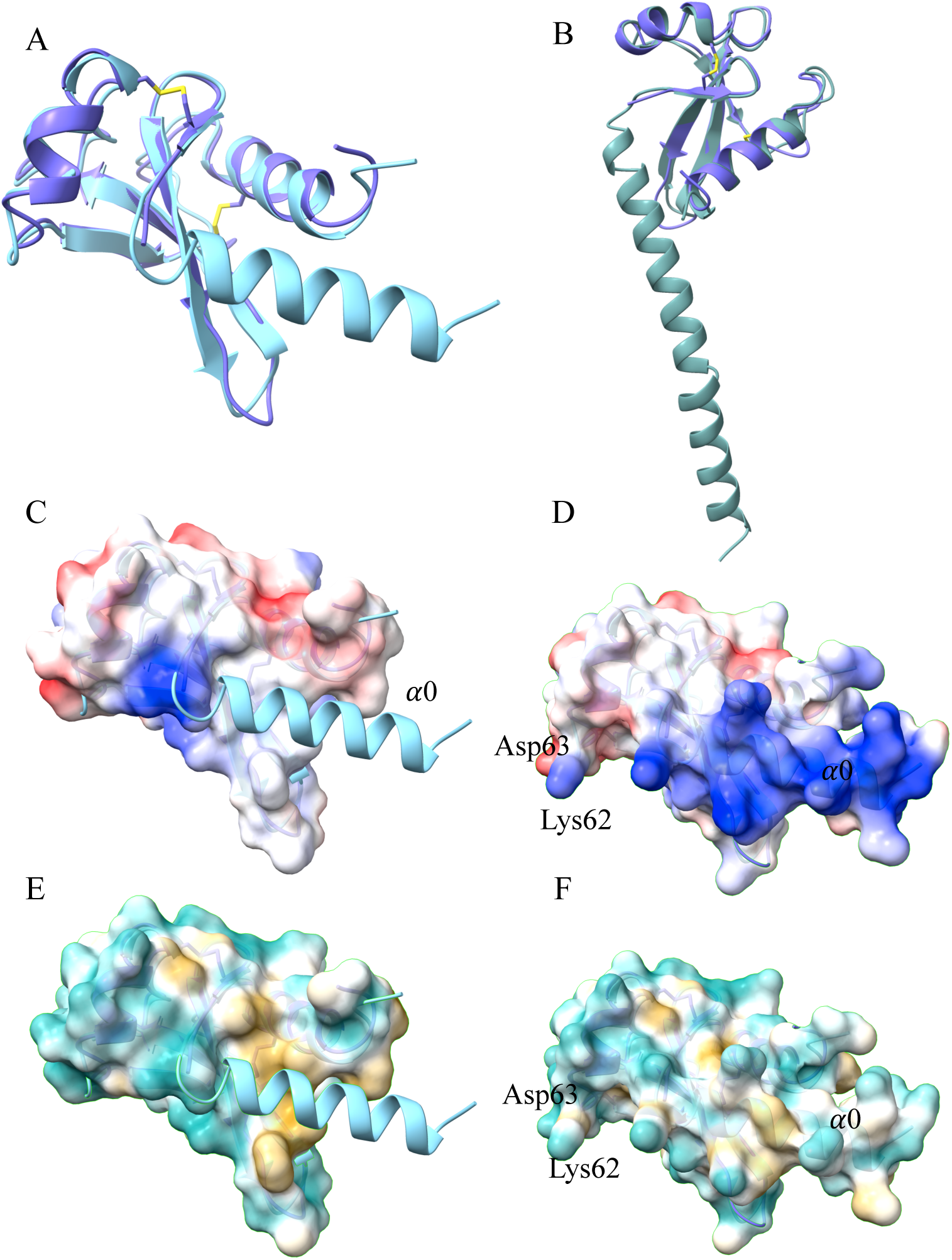
Comparison of the X-ray crystal structure with the AlphaFold 2.0 model of A21. **A**, Comparison of the A21 X-ray structure (41-117, purple) with the AlphaFold2.0 model (24-117, cyan), revealing a root mean squared deviation (RMSD) of 1.3 Å using 75 Cα pairs. Note the N-terminal α0-helix in the foreground (cyan) is modeled by AlphaFold2.0 but was disordered in the X-ray structure. **B**, Comparison of the A21 X-ray structure (41-117, purple) with the full-length A21 AlphaFold2.0 generated model (1-117, teal) that includes the predicted A21 transmembrane domain. Note that the predicted transmembrane helix extends from the α0-helix, together making a longer predicted helix. **C-D**, Comparison of electrostatic potential between surfaces of the X-ray structure (panel C) and of the AlphaFold2.0 model (panel D). In panel D, note the positively charged (blue) region arising from the modeled N-terminal α0-helix (α0), that is shown in panel C as a ribbon (cyan). **E-F**, Analogous to panels C and D, comparison of hydrophobicity between surfaces of the X-ray structure (panel E) and of the AlphaFold2.0 model (panel F). In panel E, note the hydrophobic surface (yellow) located under the modeled α0-helix (cyan ribbon).

### A21 sequence conservation in other poxviruses

Orthologs of A21 are present across all poxviruses, as shown in the phylogenetic tree (Fig. S3A). Pairwise identity with the VACV sequence varies significantly among different poxviruses (Fig. S3B). Alphaentomopoxvirus acuprea (YP_009001513.1) and Salmon gill poxvirus (SGP, CAJ1637898.1) exhibit less than 20% identity with VACV, while Yokapox (YP_004821477.1) shows the highest identity of 71% with VACV. Cervidpox, Swinepox, and Eptesipox orthologs display pairwise identities with VACV of 58-59%. The remaining sequences fall within a range of 24-54% identity with VACV. Notably, SGP represents the deepest branch of the chordopoxviruses [30]. Alignment of the amino acid sequences of the A21 orthologs from VACV and SGP indicates a strikingly low identity of only 19% (Fig. S3B). Nevertheless, AlphaFold2 predicts similar full-length structures of VACV and SGP suggesting preservation of function over hundreds of thousands of years of evolution (Fig. 3A).

**Figure 3.**
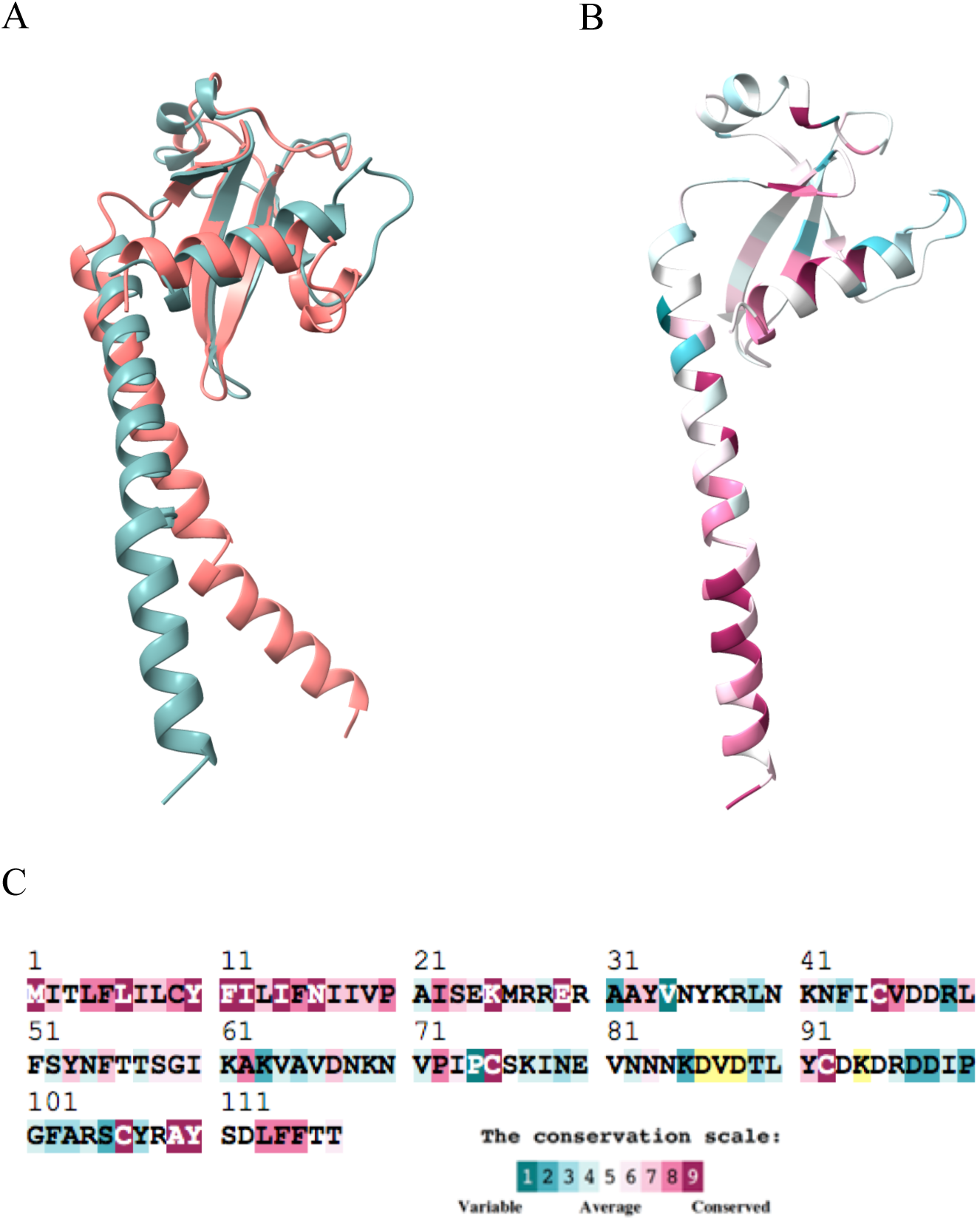
A21 sequence conservation analysis. **A**, Comparison of the AlphaFold2.0 generated models of VACV A21 (1-117, teal) with the A21 ortholog from salmon gill poxvirus (SGP) (salmon in color). The RMSD between 34 Cα pairs is 0.8 Å. **B**, VACV A21 model painted according to the degree of the protein sequence conservation among nineteen poxviruses and represented by the color scale in panel C. **C**, VACV A21 sequence color-coded to show conservation. Note the high number of conserved residues at the N-terminal and C-terminal portions of A21. Color coding is as shown in inset. Some residues had insufficient data (yellow) due to insertions and deletions.

Conservation analysis among 18 poxviruses was run on the ConSurf server [30] using the AlphaFold model with residues 1-117 as input (Fig. 3B-C). The conservation analysis revealed a high degree of conservation in the first 38 N-terminal residues, comprising the TM domain and the α0-helix, likely disordered in the crystal structure. Cys45, Cys75, Cys92, and Cys106 are conserved and preserve the fold of A21. Fig. 3 shows that many conserved amino acid residues concentrate around the hydrophobic patch formed by β1- and β2-strands, and loop 1, and α0- and α2-helix. Conserved Asp76, along with the main chain hydrogen bonding mentioned above, provides for conservation of the loop 2 shape and organization.

### Modeling of A21 interactions with other members of the EFC

Previous studies demonstrated that expression of A21 is required for assembly of the EFC [9], implying interactions with other components. Knowing the A21 structure and its agreement with the AlphaFold2 model, we set out to probe its predicted interactions with the other EFC proteins. First, we modeled its pairwise complexes with individual proteins and known sub-complexes, including O3, F9, J5, L1, A16/G9, A28/H2, and G3/L5. Except for O3, the structure of which has not been determined, the AlphaFold2 models agreed well with the experimentally determined structures. This pairwise modeling suggested that out of ten EFC proteins, A21 only interacts with G3/L5 and the small 35 amino acid O3 that consists mostly of a TM domain. To consider the arrangement of the interacting partners around A21, we modeled the entire A21, G3, L5, and O3 subcomplex and analyzed the interactions within it (Fig. 4). The analysis revealed that the N-terminal TM domain, loop 1, loop 3, and the C-terminal α-helix of A21 are the major areas of its interactions with the other elements of the subcomplex. The interface between A21 and L5 has an area of 1033 A^2^ and contains a hydrogen bond between the A21/Tyr107 hydroxyl and the main chain oxygen of L5/Met53 with a salt bridge between A21/Lys25 and L5/Glu50 (Fig. 4C). The interface between A21 and G3 is slightly smaller at 903 A^2^, with one bond between A21/Tyr10 and G3/Leu9 with a salt bridge between A21/Arg104 and G3/Glu38. The C-terminal α-helix and N-terminal TM domain of A21 provide a cluster of residues that interacts extensively with the G3/L5 subcomplex. On the other side of the A21 TM domain, Phe55 at the tip of the β2-sheet and Thr57 of loop 1 form a binding site for O3 with an interface area of 750 A^2^. The binding of O3 by A21 is stabilized by hydrogen bonds between A21/Lys37 and O3/Leu31 with a salt bridge between A21/Arg30 and O3/Glu30. To our knowledge, this is the first time that modeling suggests an interaction between A21 and O3. An interaction of A21 with O3 was previously suggested based on split GFP interactions [31]. The subsequent PISA (Protein Interfaces, Surfaces and Assemblies, Fig. 4D) analysis of interfaces of the modeled A21, G3, L5, and O3 subcomplex implies that the N-terminal TM domain of A21, along with its C-terminal α2-helix, are the major structural elements responsible for A21 integration in EFC. This notion agrees with the conservation analysis data indicating high conservation of the residues 1-30 (TM and α0-helix) and 106-117 (α2-helix) throughout A21 orthologs.

**Figure 4.**
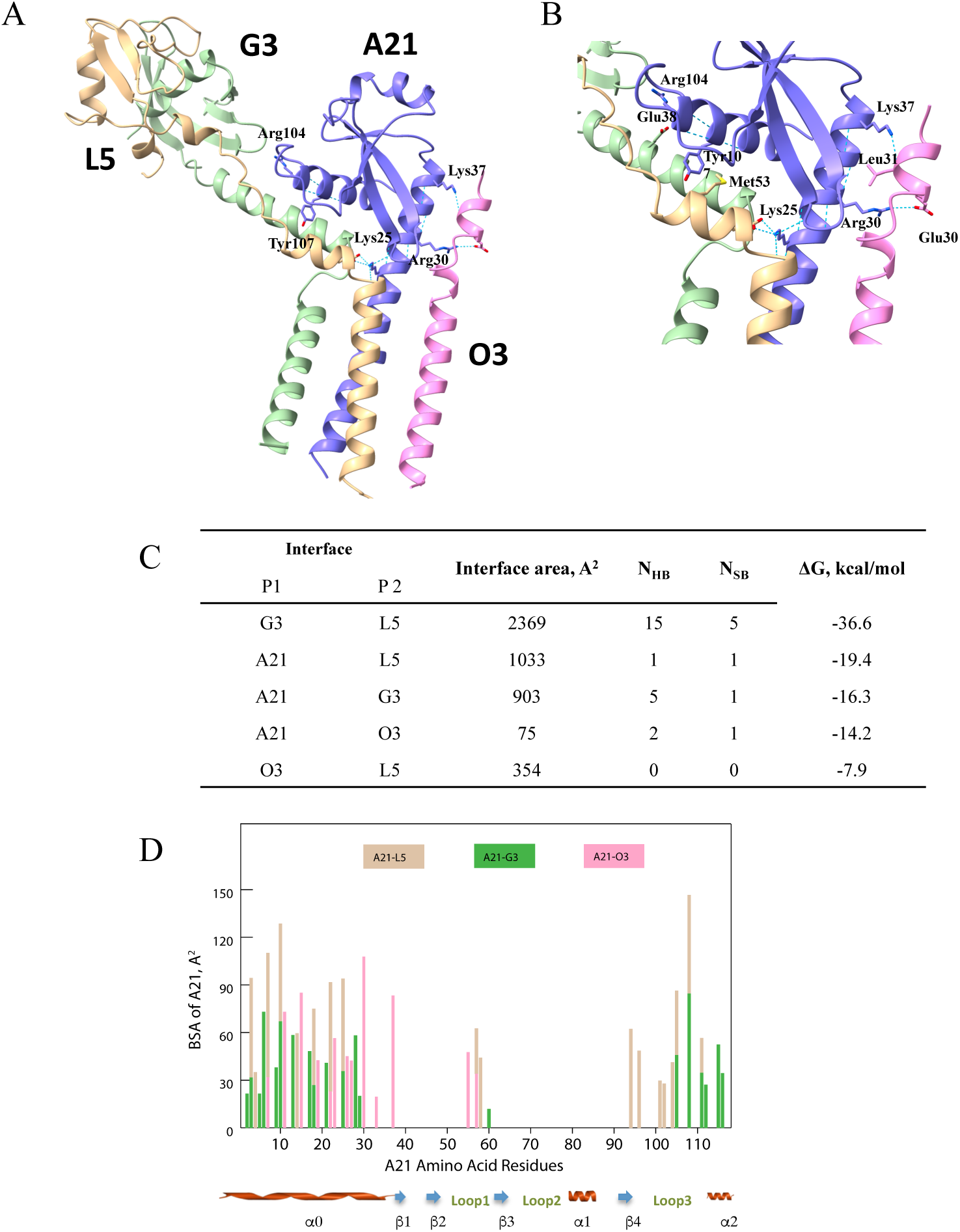
Modeling of A21/O3/G3/L5 subcomplex. **A**, Overview of the subcomplex with A21 in purple, O3 in pink, G3 in green, and L5 in beige. Key A21 residues providing for subcomplex formation are labeled. **B**, Close up of a portion of the subcomplex including the modeled A21 α0-helix making contacts with O3. Sidechains for subcomplex residues that participate in charge-charge and hydrogen bonding interactions between subunits are shown. **C**, Analysis of the subcomplex interfaces with the PISA server [46, 50] (see “Methods”). N_HB_, number of hydrogen bonds and N_SB_, number of salt bridges. **D**, Buried surface area (BSA) per A21 residue in the A21/G3/L5/O3 subcomplex. For example, at A21 residue position 30 on the x-axis, a pink bar extends to a BSA on the y-axis of about 110 Å^2^, signifying the interaction between residue 30 and the O3 subunit (pink). The BSA of some A21 residues result from interactions with two subunits. An example of this is at position 10 on the x-axis, where G3 (green) contributes about 70 Å^2^ and L5 (beige) contributes about 130 Å^2^ to the BSA of residue 10. Note that the majority of interactions and BSA occur at the N-terminal and C-terminal portions of the A21 molecule.

In conclusion, the highly conserved TM-domain and α0-helix along with α2-helix provide for integration of A21 and its orthologs into EFC, primarily interacting with G3, L5, and O3. Conservation of the overall fold of A21 and its orthologs relies on two disulfide bonds, the hydrogen bonds between the main-chain atoms, and several highly conserved residues (Phe55, Gly59, and Asp67). Hence, sequence conservation in the three loops and four β-strands of A21 provide resistance to evolutionary variation of the A21 structure. With A21, the structures of each of the EFC proteins are now known. Further extensive research is needed to elucidate the overall EFC assembly and its mechanism of action.

## Material and Methods

### Expression of A21 in *E. coli* NiCo21(DE3)

The E.coli codon-optimized DNA encoding amino acid residues 24 – 117 of VACV strain WR A21 ectodomain modified as shown in Fig. 1A with N-terminal 6xHis tag, spacer, and TEV protease cleavage site was synthesized by Blue Heron Biotech and inserted into pNAN expression vector [32]. The plasmid was used to transform NiCo21(DE3) cells (New England BioLabs) and A21Δ_N_23 was expressed as previously described [32]. Briefly, *E. coli* culture was grown to OD_600_ of 0.5 – 0.6, and the protein expression was induced with 1 mM isopropyl β-D-thiogalactoside (IPTG). The cells were harvested by centrifugation; the pellet was washed once in 50 mM Tris-HCl pH 8.0, 25% sucrose, 0.1% NaN_3_, 1 mM EDTA, and 10 mM 1,4-dithiothreitol (DTT). The cells were then resuspended in five volumes of lysis buffer (50 mM Tris-HCl pH 8.0, 1% Triton X-100, 1% Na-deoxycholate, 0.1% NaN_3_, 100 mM NaCl, 10 mM DTT, 5 mM MgCl_2_) with 1 mg/ml lysozyme and 750 U benzonase and lysed by three cycles of consecutive freeze-thawing. The IBs were harvested by centrifugation, washed, and homogenized three times in the lysis buffer without MgCl_2_. Following a triple wash in 50 mM Tris-HCl, 0.1% NaN_3_, 100 mM NaCl, 1 mM EDTA, 10 mM DTT, the IBs were denatured and dissolved in 100 mM Tris-HCl, pH 8.0, 6 M guanidine-HCl, 10 mM DTT, 1 mM EDTA overnight at 4°C. Insoluble material was removed by 1 h centrifugation at 10,000 × g and solubilized IBs were further stored at −80°C until use.

### Refolding and Purification of A21

Refolding was carried out by rapid dilution of solubilized IBs into 1 L of 50 mM Tris-HCl, pH 8.0, 50 mM NaCl, 500 mM L-arginine-HCl, 2 mM EDTA, 40 mM sucrose, 10 mM DTT, 5 mM cystamine-HCl similar to the protocol previously described [32]. After refolding, any remaining free cysteines were oxidized by adding 50 mM cystamine-HCl. The solution was dialyzed twice against 50 mM Tris-HCl, 50 mM NaCl pH 8.0. Dialyzed and refolded A21 was sterile filtered and loaded on Ni-NTA resin. The resin was washed with three column volumes of 50 mM Tris-HCl, 50 mM NaCl pH 8.0, and A21 was eluted with an increasing imidazole gradient. Fractions containing A21 were dialyzed against 50 mM Tris-HCl, 50 mM NaCl, pH 8.0 to remove imidazole. A21 concentration was measured spectrophotometrically at 280 nm using the molar extinction coefficient of 10,680 M^-1^cm^-1^. To remove the N-terminal Hisx6-tag, A21 was incubated with recombinant His-tagged TEV protease [33] (10:1 w/w A21:protease) overnight at room temperature. Uncleaved A21 and recombinant protease were removed by passing the solution over an Ni-NTA column. Cleaved A21 was further purified by SEC in 10 mM Tris-HCl, 50 mM NaCl, pH 8.0. The monomeric fraction was collected and concentrated to 10 mg/ml. Precipitated material was removed by high-speed centrifugation, and the integrity of the concentrated protein was monitored by DLS. The purity of the protein was > 95% as assessed by SDS-PAGE.

### Crystallization of A21

High-quality rod-shaped crystals were obtained by the hanging drop vapor diffusion method with 20% PEG 4000, 0.1 M Na citrate, pH 5.5 crystallization solution. The drops were set up by mixing 1 μl of 10 mg/ml protein with 1 μl of well solution and 2% w/v benzamidine-HCl. For cryo-protection and phasing, the crystals were incubated in 20% PEG 20000, 0.1 M Na citrate pH 5.5, 25% glycerol, and 1 M NaI for 1 min at room temperature, and the crystals were flash frozen in liquid nitrogen for X-ray data-collection.

### Data collection and structure determination

X-ray data on A21 crystals soaked with NaI [34–36] were collected on a RIGAKU X-ray source with MicroMax-002+ high-intensity microfocus sealed tube X-ray generator and an R-Axis IV++ image plate detector using Cu radiation. The data processed with XDS [37] showed the presence of a strong anomalous signal and were used in the AutoSol module of the Phenix software package [37, 38] for phasing by single anomalous diffraction (SAD). AutoSol located nine iodide sites, and anomalous difference maps confirmed that the space group was *P2_1_*. Automatic building of the A21 protein model was able to locate six A21 molecules per asymmetric unit, which was consistent with the prediction based on the calculation of the Matthews coefficient [39–41]. The auto-built model was 60% complete. It was further improved by employing cycles of refinement using the programs CNS [42] and Phenix followed by manual model rebuilding using the described programs [43, 44] into new electron density maps calculated after each refinement cycle until no further improvement of the model could be achieved at the 2.30 Å maximum resolution. The A21 structure and electron density maps generated in this study have been deposited in the RCSB Protein Data Bank with accession code 8U0R.

### AlphaFold2.0 modeling of A21 structure

The sequences of the full-length A21, the ectodomain of A21, and pairwise complexes of A21 with G3/L5, F9, O3, A16/G9, A28/H2, J5, and L1 were structurally modeled using the default settings of AlphaFold2. Where appropriate, we used PDB100 template mode that relies on the crystal structures available in PDB to date [45]. We performed structural analysis with the PISA (Proteins, Interfaces, Structures, and Assemblies) server [46]

### Phylogenetic tree and protein sequence conservation analysis of A21

An example sequence of A21 homolog was collected from each *Poxviridae* genus except for *Gammaentopoxvirus*, where no sequence was identified. In total, 22 sequences were retrieved and aligned in MAFFT [47] with BLOSUM30 and ‘leave gappy regions’ settings. Sequence identities and similarities were calculated with SIAS ( (longest sequence). For similarity calculation, aromatic amino acids were F, Y, and W, aliphatic amino acids were V, I, and L, positively charged amino acids were R, K, and H, and negatively charged amino acids were D and E. Small amino acids S and T and polar (not charged) amino acids N and Q were grouped. Calculation of identity and similarity was performed by equation:

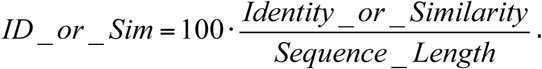

The alignment was used to generate the phylogenetic tree (Auto substitution model) using IQ-TREE [48]. The tree was calculated with 1000 bootstrap branch support.

Conservation analysis of full-length and ectodomain of A21 was carried out using the ConSurf Server with default parameters [49]. Homologs were collected from 22 sequences used for construction of the phylogenetic tree. The E-value cutoff was 0.0001, and the number of iterations was 1. CD-HIT cutoff of 95% corresponded to the maximal sequence identity between homologs. The maximum number of final homologs was 150. The maximal overlap between homologues was 10%. The highest-scoring homolog is chosen if the overlap between two homologs exceeds 10%. The homolog’s minimal percentage of the query sequence covered was 60%. Minimal sequence identity with the query sequence was 35%. Multiple Sequence Alignment was built using MAFFT. The phylogenetic tree was constructed using Neighbor Joining with ML distance. Conservation Scores were calculated using the Bayesian method. The amino acid residue substitution model was chosen as the best fit.

## Acknowledgements

Support was provided by the Division of Intramural Research, NIAID.

## Author contributions: CRediT

**Ulrike S. Diesterbeck**: Investigation, Writing - Original Draft. **Liya A. Muslinkina**: Investigation, Visualization, Writing - Original Draft, Writing - Review & Editing. **Apostolos G. Gittis**: Investigation, Formal analysis, Data Curation. **Kavita Singh**: Investigation. **Bernard Moss**: Conceptualization, Writing - Original Draft, Writing - Review & Editing. **David N. Garboczi**: Writing - Original Draft, Writing - Review & Editing, Supervision.

---

**Table S1.** A21 data collection and refinement statistics.

| **PDB ID: 8U0R** |  |
| --- | --- |
| **Data collection** |  |
| Space group | $P2_1$ |
| Cell dimensions |  |
| $a, b, c$ (Å) | 46.86, 70.06, 78.95 |
| $\alpha, \beta, \gamma$ (°) | 90.00, 95.29, 90.00 |
| Resolution (Å) | 39.31 - 2.30 (2.38 - 2.30)* |
| $R_{\text{sym}}$ or $R_{\text{merge}}$ | 0.101 (0.333) |
| $I / \sigma I$ | 8.2 (3.1) |
| Completeness (%) | 99.2 (92.2) |
| Redundancy | 3.72 (3.25) |
| **Refinement** |  |
| Resolution (Å) | 39.31 - 2.30 |
| No. reflections | 22,602 |
| $R_{\text{work}} / R_{\text{free}}$ | 0.211/0.239 |
| No. atoms |  |
| Protein | 3,389 |
| Ligand/ion | 213 |
| Water | 85 |
| $B$-factors [Å<sup>2</sup>] | |
| Protein | 50.69 |
| Ligand/ion | 56.29 |
| Water | 50.16 |
| R.m.s deviations |  |
| Bond lengths (Å) | 0.011 |
| Bond angles (°) | 1.301 |

\*Highest resolution shell is shown in parenthesis.

**Table S2.** Hydrogen bonds occurring in A21 loops.

| A21 atoms | Hydrogen bond length (Å) |
| --- | --- |
| Loop 1 |  |
| **Asn54 (N)···Lys61 (O)** * | 3.0 |
| **Asn54 (O)···Lys61 (N)** | 3.0 |
| Asn54 (OD1)···Lys63 (NZ) | 2.5 |
| Asn54 (OD1)···Tyr91 (OH) | 3.0 |
| Loop 2 |  |
| **Ala65 (O)···Ile73 (N)** | 3.2 |
| **Asp67 (N)···Val71 (O)** | 2.7 |
| **Asp67 (O)···Asn70 (N)** | 2.8 |
| Asp67 (OD1)···Lys69 (N) | 2.9 |
| Asp67 (OD1)···Val71 (N) | 2.7 |
| **Pro74 (O)···Lys77 (N)** | 3.2 |
| Loop 3 |  |
| **Asp93(O)···Arg96 (N)** | 3.1 |
| **Ile99 (O)···Phe102 (N)** | 3.3 |
| **Gly101 (O)···Ser105 (N)** | 3.0 |
| **Ala62 (O)···Cys92 (N)** | 2.8 |
| Asp93 (OD1)···Asp95 (N) | 2.6 |
| Arg96 (NH1)···Asp98 (OD1) | 3.0 |
| Arg49 (NH1)···**Pro100 (O)** | 3.3 |
| Arg49 (NH2)···**Pro100 (O)** | 2.8 |
| Ser105 (OG)···**Gly101 (O)** | 3.0 |

\*Main chain N and O hydrogen bonds are given in bold.

## Notes

### Competing Interest Statement

The authors have declared no competing interest.

## References

[1] Satheshkumar PS, Damon I. Poxviruses. In: Howley PM, Knipe DM, Cohen JL, Damania BA, editors. Fields Virology: DNA Viruses2021. p. 614–40.

[2] Moss B. Understanding the biology of monkeypox virus to prevent future outbreaks. Nature Microbiology. 2024;9:1408–16.

[3] Moss B, Smith GL. Research with variola virus after smallpox eradication: Development of a mouse model for variola virus infection. Plos Pathogens. 2021;17.

[4] Moss B. Poxvirus cell entry: how many proteins does it take? Viruses. 2012;4:688–707.

[5] Matamoros T, Alejo A, Rodriguez JM, Hernaez B, Guerra M, Fraile-Ramos A, et al. African Swine Fever Virus Protein pE199L Mediates Virus Entry by Enabling Membrane Fusion and Core Penetration. Mbio. 2020;11.

[6] Kao S, Kao CF, Chang W, Ku C. Widespread Distribution and Evolution of Poxviral Entry-Fusion Complex Proteins in Giant Viruses. Microbiol Spectr. 2023;11:e0494422.

[7] Ojeda S, Domi A, Moss B. Vaccinia virus G9 protein is an essential component of the poxvirus entry-fusion complex. J Virol. 2006;80:9822–30.

[8] Ojeda S, Senkevich TG, Moss B. Entry of vaccinia virus and cell-cell fusion require a highly conserved cysteine-rich membrane protein encoded by the A16L gene. J Virol. 2006;80:51–61.

[9] Senkevich TG, Ojeda S, Townsley A, Nelson GE, Moss B. Poxvirus multiprotein entry-fusion complex. Proc Natl Acad Sci USA. 2005;102:18572–7.

[10] Bisht H, Weisberg AS, Moss B. Vaccinia virus L1 protein is required for cell entry and membrane fusion. J Virol. 2008;82:8687–94.

[11] Brown E, Senkevich TG, Moss B. Vaccinia virus F9 virion membrane protein is required for entry but not virus assembly, in contrast to the related l1 protein. J Virol. 2006;80:9455–64.

[12] Townsley A, Senkevich TG, Moss B. The product of the vaccinia virus L5R gene is a fourth membrane protein encoded by all poxviruses that is requried for cell entry and cell-cell fusion. J Virol. 2005;79:10988–98.

[13] Satheshkumar PS, Moss B. Characterization of a newly Identified 35 amino acid component of the vaccinia virus entry/fusion complex conserved in all chordopoxviruses. J Virol. 2009;83:12822–32.

[14] Senkevich TG, Moss B. Vaccinia virus H2 protein is an essential component of a complex involved in virus entry and cell-cell fusion. J Virol. 2005;79:4744–54.

[15] Senkevich TG, Ward BM, Moss B. Vaccinia virus entry into cells is dependent on a virion surface protein encoded by the A28L gene. J Virol. 2004;78:2357–66.

[16] Moss B. Membrane fusion during poxvirus entry. Semin Cell Dev Biol. 2016;60:89–96.

[17] Senkevich TG, White CL, Koonin EV, Moss B. Complete pathway for protein disulfide bond formation encoded by poxviruses. Proc Natl Acad Sci USA. 2002;99:6667–72.

[18] Laliberte JP, Weisberg AS, Moss B. The membrane fusion step of vaccinia virus entry is cooperatively mediated by multiple viral proteins and host cell components. PLoS Pathog. 2011;7:e1002446.

[19] Nelson GE, Wagenaar TR, Moss B. A conserved sequence within the H2 subunit of the vaccinia virus entry/fusion complex is important for interaction with the A28 subunit and infectivity. J Virol. 2008;82:6244–50.

[20] Wagenaar TR, Ojeda S, Moss B. Vaccinia virus A56/K2 fusion regulatory protein interacts with the A16 and G9 subunits of the entry fusion complex. J Virol. 2008;82:5153–60.

[21] Wolfe CL, Moss B. Interaction between the G3 and L5 proteins of the vaccinia virus entry-fusion complex. Virology. 2011;412:278–83.

[22] Su HP, Garman SC, Allison TJ, Fogg C, Moss B, Garboczi DN. The 1.51-A structure of the poxvirus L1 protein, a target of potent neutralizing antibodies. Proc Natl Acad Sci USA. 2005;102:4240–5.

[23] Diesterbeck US, Gittis AG, Garboczi DN, Moss B. The 2.1 angstrom structure of protein F9 and its comparison to L1, two components of the conserved poxvirus entry-fusion complex. Scientific Reports. 2018;8.

[24] Lin S, Yue D, Yang FL, Chen ZM, He B, Cao Y, et al. Crystal structure of vaccinia virus G3/L5 sub-complex reveals a novel fold with extended inter-molecule interactions conserved among orthopoxviruses. Emerging Microbes & Infections. 2023;12.

[25] Yang F, Lin S, Chen Z, Yue D, Yang M, He B, et al. Structural basis of poxvirus A16/G9 binding for sub-complex formation. Emerg Microbes Infect. 2023;12:2179351.

[26] Kao CF, Tsai MH, Carillo KJ, Tzou DL, Chang W. Structural and functional analysis of vaccinia viral fusion complex component protein A28 through NMR and molecular dynamic simulations. PLoS Pathog. 2023;19:e1011500.

[27] Kao CF, Liu CY, Hsieh CL, Carillo KJD, Tzou DL, Wang HC, et al. Structural and functional analyses of viral H2 protein of the vaccinia virus entry fusion complex. J Virol. 2023;97:e0134323.

[28] Carillo KJ, Tzou DL. Vaccinia virus J5 ectodomain. PDB2023.

[29] Townsley A, Senkevich TG, Moss B. Vaccinia virus A21 virion membrane protein is required for cell entry and fusion. J Virol. 2005;79:9458–69.

[30] Yariv B, Yariv E, Kessel A, Masrati G, Chorin AB, Martz E, et al. Using evolutionary data to make sense of macromolecules with a “face-lifted” ConSurf. Protein Sci. 2023;32:e4582.

[31] Schin AM, Diesterbeck US, Moss B. Insights into the Organization of the Poxvirus Multicomponent Entry-Fusion Complex from Proximity Analyses in Living Infected Cells. J Virol. 2021;95:e0085221.

[32] Su HP, Lin DYW, Garboczi DN. The structure of G4, the poxvirus disulfide oxidoreductase essential for virus maturation and infectivity. Journal of Virology. 2006;80:7706–13.

[33] Tropea JE, Cherry S, Waugh DS. Expression and purification of soluble His(6)-tagged TEV protease. Methods Mol Biol. 2009;498:297–307.

[34] Dauter Z, Dauter M. Entering a new phase: using solvent halide ions in protein structure determination. Structure. 2001;9:R21–6.

[35] Douglas AE, Corbett KD, Berger JM, McFadden G, Handel TM. Structure of M11L: A myxoma virus structural homolog of the apoptosis inhibitor, Bcl-2. Protein Sci. 2007;16:695–703.

[36] Nagem RA, Polikarpov I, Dauter Z. Phasing on rapidly soaked ions. Methods Enzymol. 2003;374:120–37.

[37] Kabsch W. Xds. Acta Crystallogr D Biol Crystallogr. 2010;66:125–32.

[38] Adams PD, Afonine PV, Bunkoczi G, Chen VB, Davis IW, Echols N, et al. PHENIX: a comprehensive Python-based system for macromolecular structure solution. Acta Crystallogr D Biol Crystallogr. 2010;66:213–21.

[39] Weichenberger CX, Rupp B. Ten years of probabilistic estimates of biocrystal solvent content: new insights via nonparametric kernel density estimate. Acta Crystallogr D Biol Crystallogr. 2014;70:1579–88.

[40] Kantardjieff KA, Rupp B. Matthews coefficient probabilities: Improved estimates for unit cell contents of proteins, DNA, and protein-nucleic acid complex crystals. Protein Sci. 2003;12:1865–71.

[41] Matthews BW. Solvent content of protein crystals. J Mol Biol. 1968;33:491–7.

[42] Brunger AT. Version 1.2 of the Crystallography and NMR system. Nat Protoc. 2007;2:2728–33.

[43] Jones TA, Zou JY, Cowan SW, Kjeldgaard M. Improved methods for building protein models in electron density maps and the location of errors in these models. Acta Crystallogr A. 1991;47 (Pt 2):110–9.

[44] Emsley P, Lohkamp B, Scott WG, Cowtan K. Features and development of Coot. Acta Crystallogr D Biol Crystallogr. 2010;66:486–501.

[45] Jumper J, Evans R, Pritzel A, Green T, Figurnov M, Ronneberger O, et al. Highly accurate protein structure prediction with AlphaFold. Nature. 2021;596:583–9.

[46] Krissinel E, Henrick K. Inference of macromolecular assemblies from crystalline state. J Mol Biol. 2007;372:774–97.

[47] Katoh K, Misawa K, Kuma K, Miyata T. MAFFT: a novel method for rapid multiple sequence alignment based on fast Fourier transform. Nucleic Acids Res. 2002;30:3059–66.

[48] Nguyen LT, Schmidt HA, von Haeseler A, Minh BQ. IQ-TREE: a fast and effective stochastic algorithm for estimating maximum-likelihood phylogenies. Mol Biol Evol. 2015;32:268–74.

[49] Landau M, Mayrose I, Rosenberg Y, Glaser F, Martz E, Pupko T, et al. ConSurf 2005: the projection of evolutionary conservation scores of residues on protein structures. Nucleic Acids Res. 2005;33:W299–302.

[50] Krissinel E. Crystal contacts as nature’s docking solutions. J Comput Chem. 2009;31:133–43.

