## Supplementary Figures for "The 2.3 Å Structure of A21, a Protein Component of the Conserved Poxvirus Entry-Fusion Complex": SUPPLEMENTARY FIGURES.pdf

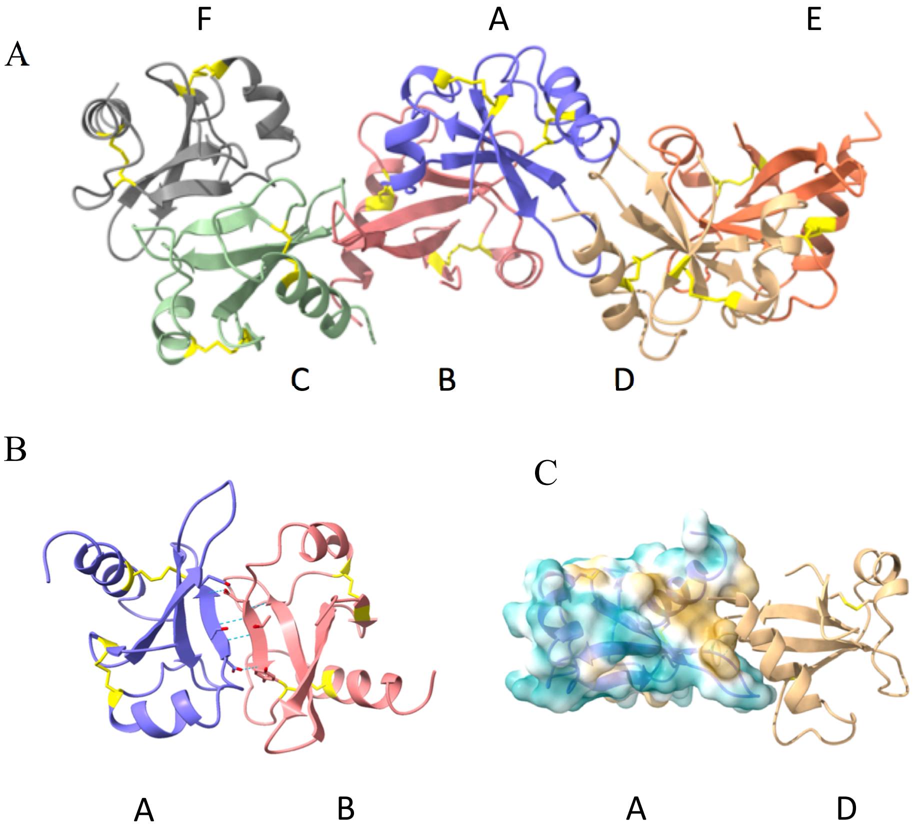


**Figure S1**.

**Figure S1.** X-ray crystal structure of A21. **A**, The six A21 molecules (A-F) that make up the asymmetric unit. **B**, The A/B interface is stabilized by two hydrogen bonds between A:Tyr91/B:Asp88 and two hydrogen bonds between A:Thr89/B:Thr89. **C**, Dispersive interactions stabilize the A/D interface. The surface of A is colored according to hydrophobicity and reveals a hydrophobic patch (yellow) in the interface with molecule D (beige ribbon). Two hydrogen bonds between A:Ile61 and D:Ile60 are present, but not shown. Disulfide bonds are shown in yellow.

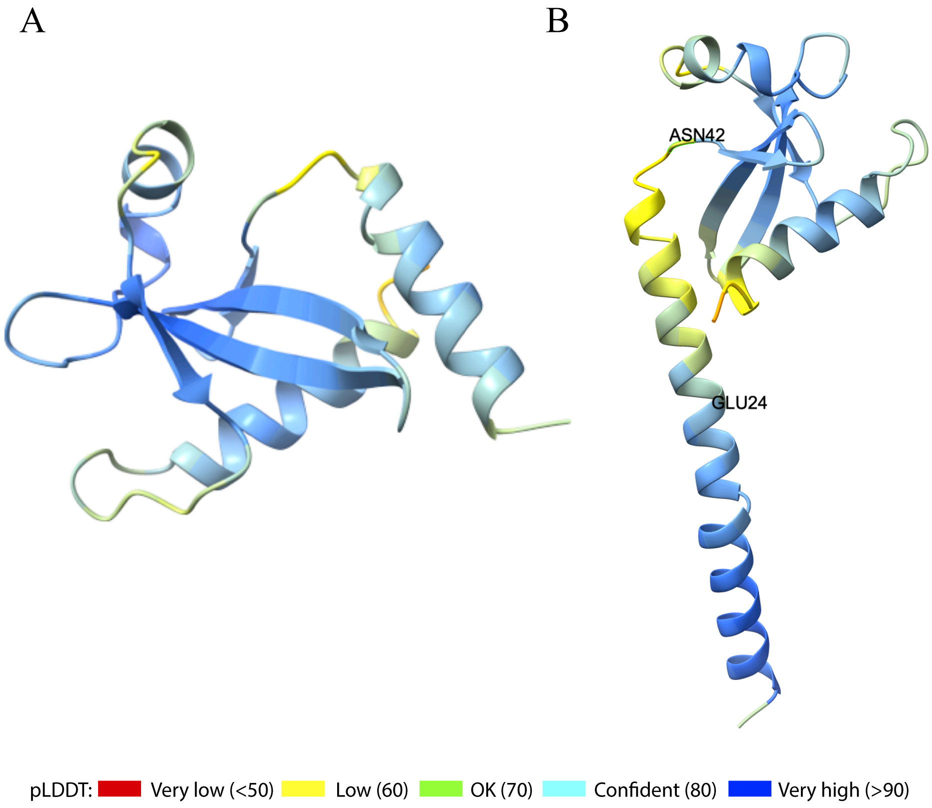
A
B
ASN42
GLU24
pLDDT: Very low (<50) Low (60) OK (70) Confident (80) Very high (>90)

**Figure S2**.

**Figure S2**. Confidence metrics for the AlphaFold2.0 models of A21 colored by predicted local distance difference test (pLDDT). **A**, Ectodomain comprising residues 24-117. **B**, Full-length A21. Note overall high confidence of prediction for A21 structure, with low confidence regions (yellow) in the α0-helix (residues Glu24-Asn42) and at the A21 C-terminus (end of α2-helix).

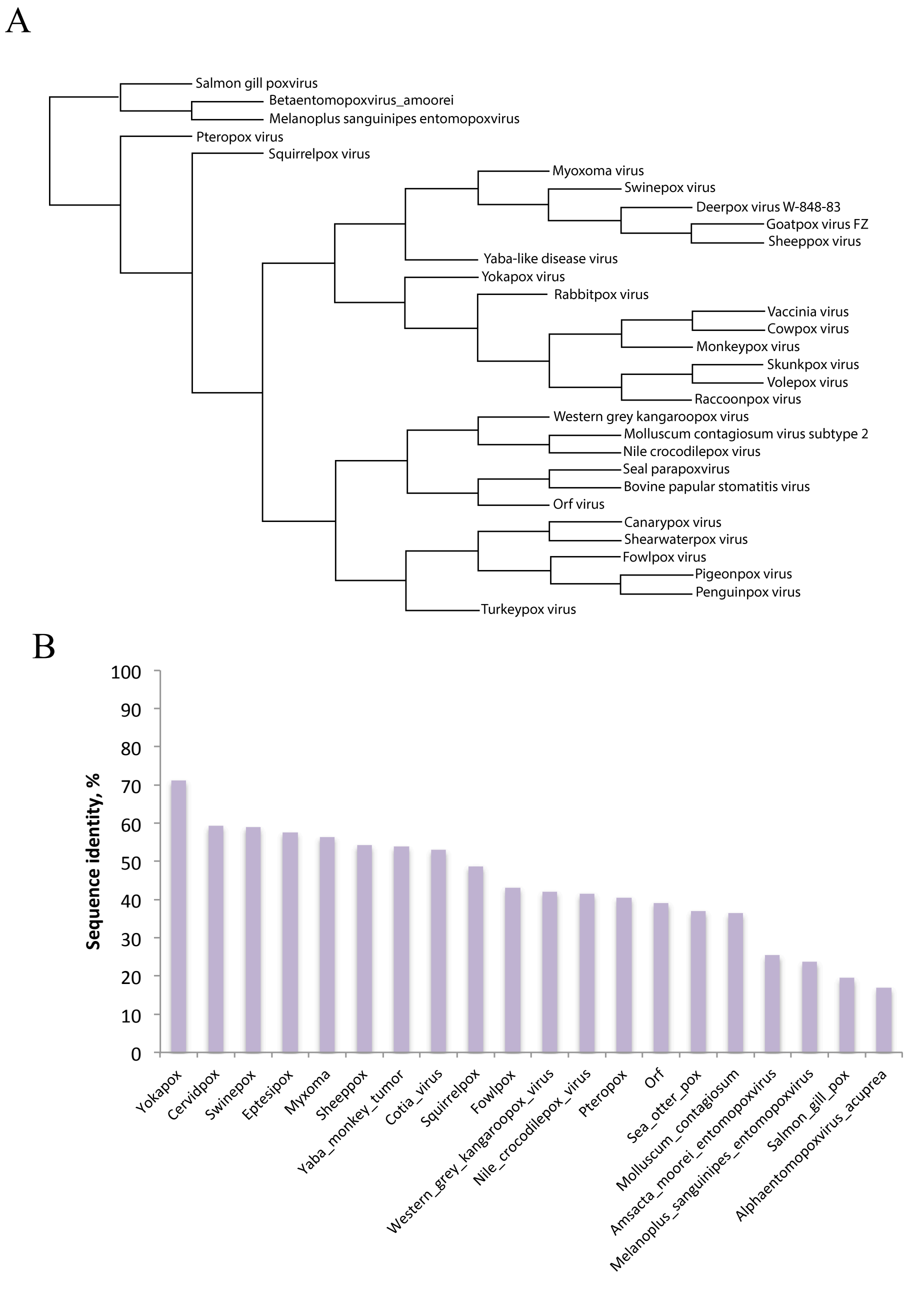

A
Salmon gill poxvirus
Betaentomopoxvirus_amoorei
Melanoplus sanguinipes entomopoxvirus
Pteropox virus
Squirrelpox virus
Myoxoma virus
Swinepox virus
Deerpox virus W-848-83
Goatpox virus FZ
Sheeppox virus
Yaba-like disease virus
Yokapox virus
Rabbitpox virus
Vaccinia virus
Cowpox virus
Monkeypox virus
Skunkpox virus
Volepox virus
Raccoonpox virus
Western grey kangaroopox virus
Molluscum contagiosum virus subtype 2
Nile crocodilepox virus
Seal parapoxvirus
Bovine papular stomatitis virus
Orf virus
Canarypox virus
Shearwaterpox virus
Fowlpox virus
Pigeonpox virus
Penguinpox virus
Turkeypox virus
B
Sequence identity, %
100
90
80
70
60
50
40
30
20
10
0
Yokapox
Cervidpox
Swinepox
Eptesipox
Myxoma
Sheeppox
Yaba_monkey_tumor
Cotia_virus
Squirrelpox
Fowlpox
Western_grey_kangaroopox_virus
Nile_crocodilepox_virus
Pteropox
Orf
Sea_otter_pox
Molluscum_contagiosum
Amsacta_moorei_entomopoxvirus
Melanoplus_sanguinipes_entomopoxvirus
Salmon_gill_pox
Alphaentomopoxvirus_acuprea


**Figure S3**.

**Figure S3**. Phylogenetic analysis of A21. **A**, Phylogenetic tree of 22 orthologs of VACV A21aligned in MAFFT [1] with BLOSUM30 and 'leave gappy regions' settings. The alignment was used to generate the phylogenetic tree using auto substitution model IQ-TREE [2]. The tree was calculated with 1000 bootstrap branch support. **B**, Pairwise sequence identity of selected A21 orthologs relative to the sequence of VACV.

A B C

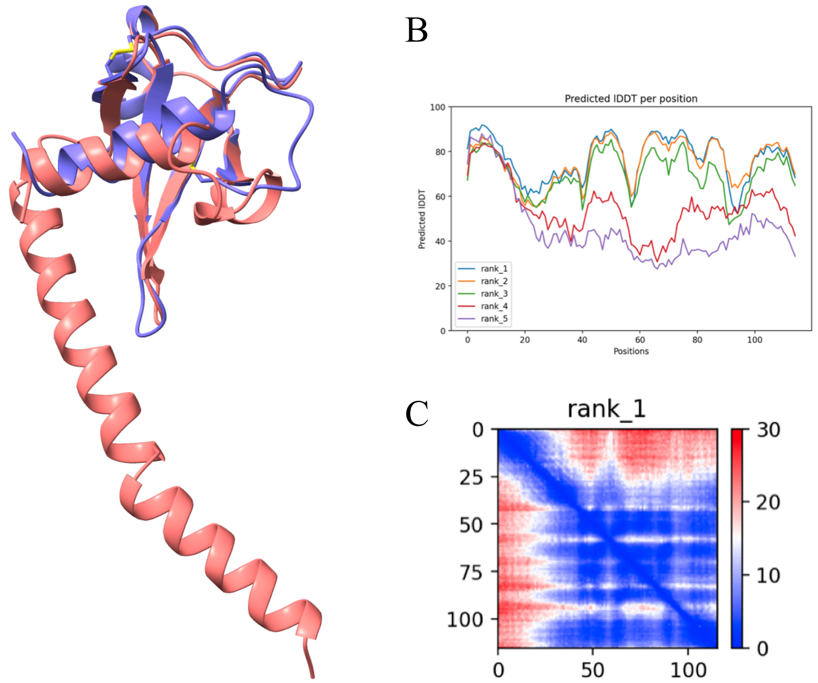


D

```
                     10        20        30        40        50        60
                      |         |         |         |         |         |
VACV WR      MITLFLILCYFILIFNIIVPAISEKMRRERAAYVNY-KRLNKNFICVDDRLFSYNFTTSG
SGP          MILFALILFFHIFLYNFVIPGVMLKKKRNNRIFNDLSTKLPSNYICINNKWATVTFDSLD
             ** : *** ::*:::*:::*::  * :*::  : :  ::* :*:**::::  : :* : :

                     70        80        90       100       110
                      |         |         |         |         |
VACV WR      IKAKVAVDNKNVPIPCSKINEVNNNKDVDTLYCDKDRDDIPGFARSCYRAYSDLFFTT-
SGP          IDVLFIKTTNGANVNCDAD--PS-NKNFTMTSCLS-TDGFALPNGECGKALKHILVSYL
             *:: :   :::: : *:     : **::    * :  *:::    :* :* ::::::
```

**Figure S4.**

**Figure S4**. Comparison of the AlphaFold2.0 generated model for a full-length salmon gill poxvirus (SGP) A21 with VACV WR A21. **A**, Overlay with x-ray crystal structure of VACV WR A21. **B**, Predicted local distance difference test (pLDDT) score per position for the highest-ranked five models generated by AlphaFold2.0. The pLDDT is plotted against the amino acid residue position. pLDDT values greater than 90 indicate high accuracy, equivalent to structures determined by experiments. Values between 70 and 90 indicate a high accuracy, where the prediction of the main chain of the protein is reliable. Values between 50 to 70 indicate a lower accuracy, but it is likely that the predictions of individual secondary structures are correct. **C**, Predicted aligned error (PAE) plot for the best model, rank_1. PAE is defined as the expected positional error (in Å) at residue X, measured if the predicted and true structures were aligned on residue Y. Both axes are labeled with the position numbers of the A21 sequence. The predicted error in position per pair of residues is color coded from blue (low error, 0 Å) to red (high error, 30 Å), as shown in the key (at right). For example, the intersection of residues 50 and 75 is blue, indicating confidence in the positions of both residues relative to each other. On the other hand, the first 30 residues are red, reflecting a high positional error against all other A21 residues. This means that the positions of the first 30 residues are not known accurately (red) relative to the rest of the molecule and is consistent with the inherent flexibility of the predicted transmembrane domain. The diagonal line (blue) shows the low error of each residue aligned on itself. **D**, Comparison of A21 protein sequences from VACV WR and SGP. Shown are 22 residues (19%) that are identical (*), 49 residues (41%) that are similar (:), and 47 residues (40%) that are different.

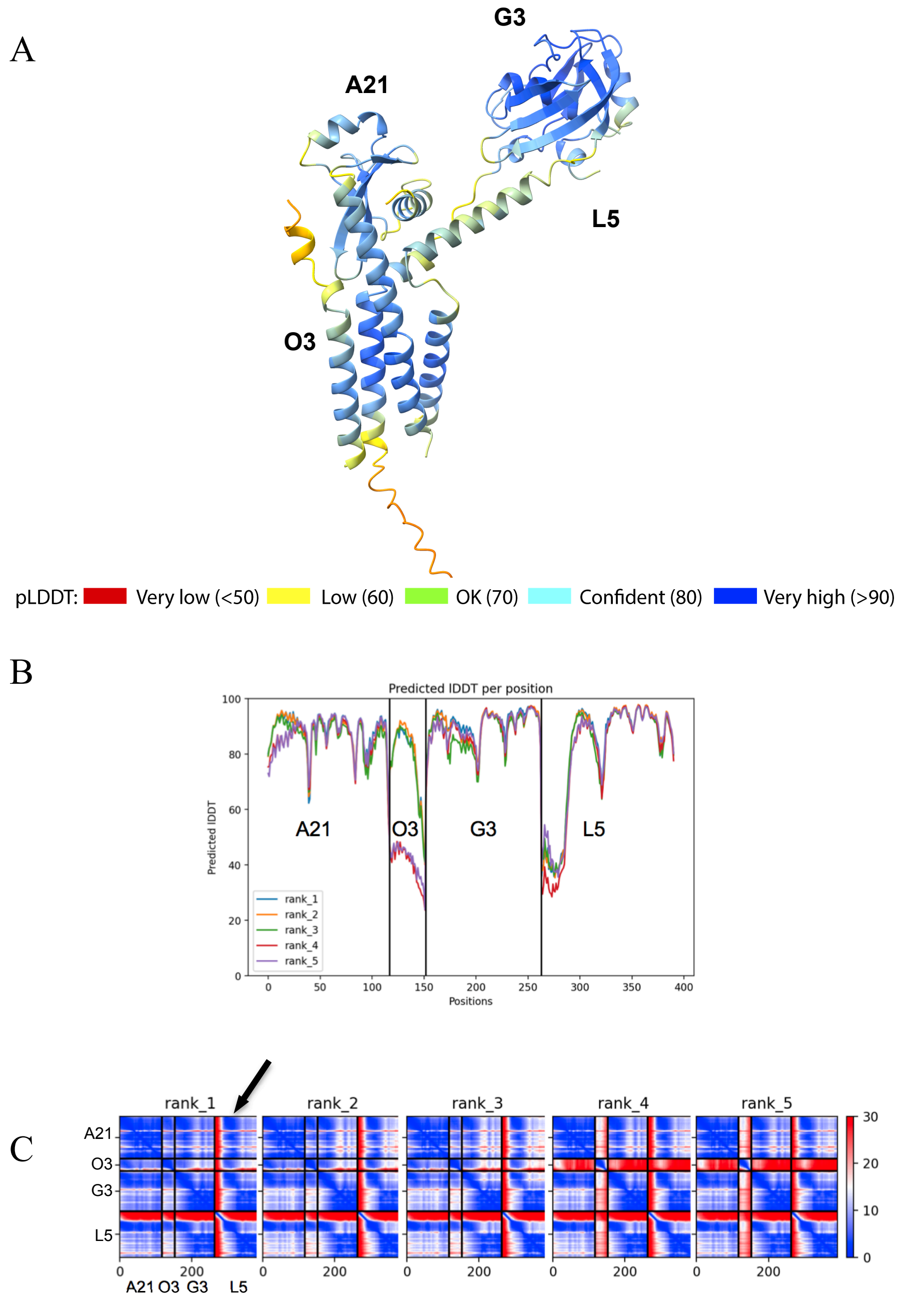


### Figure S5.

**Figure S5**. Confidence metrics for the predicted structure of the A21/O3/G3/L5 subcomplex. **A**, A21/O3/G3/L5 subcomplex structure colored by pLDDT. **B**, Predicted local distance difference test (pLDDT) score per position for the five models (rank_1 to rank_5) generated by AlphaFold2.0. See legend to Fig. S4B for description of pLDDT. **C**, Prediction aligned error (PAE) plots for models rank_1 to rank_5. See legend to Fig. S4C for description of PAE. At upper left in rank_1, A21 is plotted against itself and is mostly blue, indicating the low error predicted in the position of A21 residues with respect to each other. At upper right in rank_1, A21 is plotted against L5 residues revealing a blue stripe of low error (arrow). The error in the position of residues within the L5 intravirion domain (N-terminal 20 residues) is high (red) for all models. The x-axes are numbered from 1 to about 400 as the AlphaFold2.0 output labels the axes as though the sequences were concatenated.

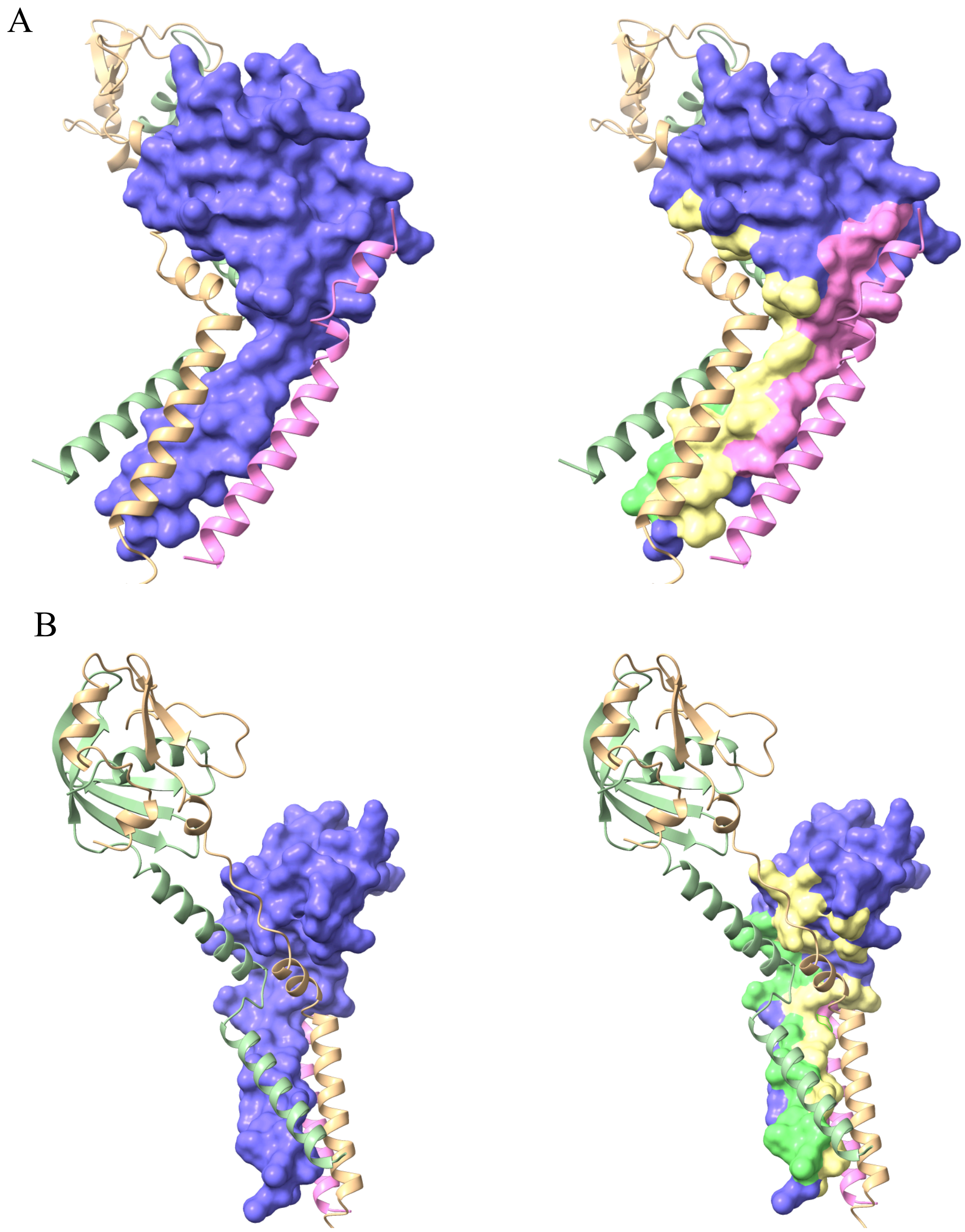


**Figure S6**.

**Figure S6**. A21/O3/G3/L5 subcomplex model showing the interactions of A21 with the rest of the complex. **A**, At left, the A21 solvent excluded surface (purple) is shown surrounded by the ribbon depictions of O3 (pink), G3 (green), and L5 (beige). At right, surfaces of A21 residues are re-colored to show which subunit (O3, G3, L5) interacts with that area of A21. Domains of G3 and L5 farther away from the transmembrane domains have been omitted for clarity. **B**, The views of the complex from panel A have been rotated to show A21 interaction surfaces with G3 and L5.

[1] Katoh K, Misawa K, Kuma K, Miyata T. MAFFT: a novel method for rapid multiple sequence alignment based on fast Fourier transform. Nucleic Acids Res. 2002;30:3059-66.

[2] Nguyen LT, Schmidt HA, von Haeseler A, Minh BQ. IQ-TREE: a fast and effective stochastic algorithm for estimating maximum-likelihood phylogenies. Mol Biol Evol. 2015;32:268-74.
